# Epigenetic features improve TALE target prediction

**DOI:** 10.1101/2021.06.14.448323

**Authors:** Annett Erkes, Stefanie Mücke, Maik Reschke, Jens Boch, Jan Grau

**Affiliations:** Institute of Computer Science, Martin Luther University Halle-Wittenberg, Halle, 06120, Germany; Department of Plant Biotechnology, Leibniz Universität Hannover, Hannover, Germany

## Abstract

The yield of many crop plants can be substantially reduced by plant-pathogenic *Xanthomonas* bacteria. The infection strategy of many *Xanthomonas* strains is based on transcription activator-like effectors (TALEs), which are secreted into the host cells and act as transcriptional activators of plant genes that are beneficial for the bacteria.

The modular DNA binding domain of TALEs contains tandem repeats, each comprising two hyper-variable amino acids. These repeat-variable diresidues (RVDs) bind to a continuous DNA stretch (a target box) and determine the specificity of a TALE. All available tools for the prediction of TALE targets within the host plant suffer from many false positives. In this paper we propose a strategy to improve prediction accuracy by considering the epigenetic state of the host plant genome in the region of the target box. To this end, we extend our previously published tool PrediTALE by two epigenetic features: (i) We allow for filtering target boxes according to chromatin accessibility and (ii) we allow for considering the methylation state of cytosines within the target box during prediction, since DNA methylation may affect the binding specificity of RVDs. Here, we determine the epigenetic features from publicly available DNase-seq, ATAC-seq, and WGBS-seq data in rice.

We benchmark the utility of both epigenetic features separately and in combination, deriving ground-truth from RNA-seq infections studies in rice. We find an improvement for each individual epigenetic feature, but especially the combination of both. Having established an advantage in TALE target predicting considering epigenetic features, we use these data for promoterome and genome-wide scans by our new tool EpiTALE, leading to several novel putative virulence targets.

Our results suggest that it would be worthwhile to collect condition-specific chromatin accessibility data and methylation information when studying putative virulence targets of *Xanthomonas* TALEs.

## INTRODUCTION

The cultivation of crop plants can be severely impaired by the infestation with phytopathogenic *Xanthomonas* bacteria. In many parts of the world, the crop yield of rice plays a key role in ensuring nutrition of the population. However, the yield of a rice field can be substantially reduced due to infection with *Xanthomonas oryzae* pv. *oryzae* (*Xoo*) or *Xanthomonas oryzae* pv.*oryzicola* (*Xoc*), which cause significant loss in many cultivation areas (1).

Host plant infection depends on the bacterial type III secretion system. Specific bacterial effector proteins are secreted into the plant cell, where they modulate plant response. Of these, transcription activator-like effectors (TALEs) are sequence-specific DNA-binding proteins that bind to host promoters to activate the expression of downstream genes. If such genes promote disease, they are termed susceptibility genes (2, 3).

TALE proteins comprise a nuclear localization signal, a modular DNA-binding domain, and an activation domain. The DNA-binding domain of natural TALEs is composed of 1.5 to 33.5 consecutive repeats, where each repeat binds to one nucleotide of the target box. Each repeat comprises ∼ 34, highly conserved, amino acids (AAs). Only the residues at position 12 and 13 are hyper-variable and are called repeat-variable diresidue (RVD). Only the second residue of the RVD binds to the target base, while the first residue has a stabilizing effect (4, 5). The surrounding conserved AAs form two alpha helices and present the RVD in a short loop to the DNA (4, 5). The C-terminal repeat is shorter then the others and is therefore called a “half repeat”.

The target boxes of a TALE can be predicted based on the one-to-one correspondence between RVD and target base (3, 6). For example, the RVD HD (His and Asp) prefers to bind to base ‘C’. Furthermore, TALE target boxes show an additional preference for the base at “position 0” directly preceding the nucleotides bound by the repeat array, which is usually ‘T’ (3, 7). As a rare exception, individual aberrant repeats of unusual length may loop out of the repeat array to allow binding to a target DNA sequence that is one bp shorter (8).

Several tools to identify potential target boxes based on the RVD sequence exists. These include the “Target Finder” of TALE-NT suite (http://tale-nt.cac.cornell.edu/) (9, 10), the tool Talvez (http://bioinfo-web.mpl.ird.fr/cgi-bin2/talvez/talvez.cgi) (11) and TALgetter (http://www.jstacs.de/index.php/TALgetter) (12). Our recently published tool PrediTALE (http://www.jstacs.de/index.php/PrediTALE) (13) models binding specificities based on quantitative data and includes further aspects of the binding of TALEs to their target boxes. It considers putative dependencies between adjacent RVDs and dependencies between the first RVD and the preference at position 0 of the target box, as well as the frame shift that may occur for aberrant repeats (8).

Prediction of TALE targets with PrediTALE achieves an improved prediction accuracy compared with previous approaches. Still, the predictions of all tools suffer from many false positives.

Hence, we propose two extensions of PrediTALE considering epigenetic features to reduce the amount of false positive predictions. First, we extend PrediTALE to consider DNA methylation information when making predictions and, second, we filter predictions using accessibility data such as DNase-seq and ATAC-seq. Our new application suite Epi-TALE contains all tools necessary for TALE target prediction incorporating epigenetic features of the target site.

DNA methylation is an epigenetic mechanism, where a methyl group is added to cytosine to form 5-methylcytosine (5mC) (14). As shown recently (15, 16), methylation alters the preference of RVDs for cytosines, which has been neglected by previous approaches for TALE target prediction. Biochemical analyses (15) showed that methylated C is bound by NG rather than HD. The RVD NG binds specifically to base T, which is structurally identical to 5mC in the part that faces the major grove of the DNA which essentially is bound by the RVD (17). Hence, the RVD NG may also bind well to 5mC in addition to T. The RVD N*, where ‘*’ represents the deletion of the 13th amino acid, is known to preferentially bind to T or C, and has been shown to also bind to 5mC and 5hmC (18).

Our aim is to improve predictions of PrediTALE by approximating the specificities of the different RVD types for methylated cytosine based on experimental data. Users of our new suite EpiTALE may then provide methylation data in addition to genomic sequence or extracted promoters, which will be considered in prediction scoring. EpiTALE is the first approach that accounts for methylated cytosine when predicting TALE target boxes.

In addition, it has been shown that the accessibility of chromatin in the area of the target site has an impact on the binding ability of TALEs (17, 19). Hence, we annotate the chromatin accessibility of predicted target sites using DNase-seq and ATAC-seq data to the predictions of EpiTALE and suggest criteria to filter putatively inaccessible target boxes.

We benchmark EpiTALE based on RNA-seq data after *Xanthomonas* infection of rice plants. Here, we consider infection studies for 3 *Xoo* and 10 *Xoc* strains, where each strain expresses a different repertoire of TALEs, with up to 27 TALEs per strain (20–23).

We further apply EpiTALE using both, methylation information and a filter based on chromatin accessibility, for genome-wide predictions and identify previously neglected putative TALE target boxes, which show a transcription response in infection experiments according to RNA-seq data.

## METHODS

### Data

#### Bisulfite sequencing data of rice

We obtained publicly available whole genome bisulfite sequencing (WGBS-seq) data of rice from the European Nucleotide Archive (ENA) https://www.ebi.ac.uk/ena available under run accessions SRR3485276 (replicate 1) and SRR3485277 (replicate 2). These data have been collected as part of a study by Zheng et al. (24), who investigated epigenetic changes under drought stress. The two WGBS-seq runs we consider in this study to determine DNA methylation levels in rice correspond to two biological replicates of Huhan3 (O. *sativa* L. ssp. *japonica*) under normal conditions.

We adapter clipped and quality trimmed these paired end reads using Trimmomatic (v0.33) (25) with parameters “CROP:80 SLIDINGWINDOW:4:28 MINLEN:20”. We mapped the processed reads to the rice genome (MSU7, http://rice.plantbiology.msu.edu/pub/data/Eukaryotic_Projects/o_sativa/annotation_dbs/pseudomolecules/version_7.0/all.dir/all.chrs.con) via Bismark (v0.20.0) (26) and Bowtie2 (v2.3.4.3) (27). We used the deduplication tool from Bismark to remove PCR artefacts.

We used the Bismark methylation extractor to determine the methylation levels and set the following parameters “– bedGraph –CX -p”. In plants, cytosine methylation occurs in the following three contexts: ‘CpG’, ‘CpHpG’ and ‘CpHpH’ (H = ‘A’, ‘C’ or ‘T’). With option “–CX”, the output contains the methylation of cytosines in all three contexts. The output contains a coverage file, which contains the columns: chromosome, start position, end position, methylation percentage, count methylated and count unmethylated. We finally merged the coverage files of both replicates by summing the counts at identical positions and then updating the methylation level. To obtain conservative methylation calls, we introduced a bias towards unmethylated cytosines in sparsely covered regions by adding a pseudo count of 1 to the count values for unmethylated cytosines.

Supplementary Figure S1 has been generated by the methylation report of ViewBS (28), and shows the distribution of methylation levels and the global methylation level of the three methylation contexts.

#### RNA-seq data

To benchmark EpiTALE, we used RNAseq data as described previously (13). Briefly, we used in-house RNA-seq infection studies of rice leaves with *Xoo* strains PXO83, PXO142, ICMP 3125^T^ and publicly available data from infection studies with *Xoc* strains BLS256, BLS279, CFBP2286, B8-12, L8, RS105, BXOR1, CFBP7331, CFBP7341, CFBP7342 (22). Genes that are differentially expressed in the infection studies compared to mock control and whose promoters contain a putative target box of a TALE are defined as true positive targets. A direct assignment to a single TALE of a strain is not possible based on the RNA-seq data, since the entire TALE repertoire of a strain acts simultaneously in the infection studies.

#### DNase-seq and ATAC-seq data

To identify accessible regions, we mapped publicly available DNase-seq and ATAC-seq data to the rice genome (MSU7). We downloaded DNase-seq reads of rice seedlings (29) from NCBI Sequence Read Archive (SRA), accession SRX038423, and used Cutadapt (30) for adapter clipping and Trimmomatic (v0.33) (25) with parameters “SLIDINGWINDOW:4:20 MINLEN:20” for quality trimming. We mapped the reads to the rice MSU7 genome using Bowtie2 (27). In the following we will refer to this DNase-seq dataset as ‘DNase’ to improve readability.

Two ATAC-seq datasets for wildtype rice are publicly available. In the first study, nucleosome-free chromatin was measured in a time series under different stress conditions (31). As we are interested in normal conditions, we only consider the control experiments from the corresponding ENA archive (accession: PRJNA305365). We refer to this dataset as ‘ATAC1’.

For the second study (32), ATAC-seq data of rice nuclei from leaf tissue where downloaded from ENA (accession: PRJNA391551) and we refer to this dataset as ‘ATAC2’.

For both datasets we used Trimmomatic (v0.39) in paired end mode for adapter clipping and trimming with parameters “ILLUMINACLIP:NexteraPE-PE.fa:2:30:10 SLIDING-WINDOW:4:20 MINLEN:20”, mapped the resulting reads with Bowtie2 to rice genome (MSU7) and removed duplicates with Samtools (33).

The mapping statistics of all three datasets are summarized in Supplementary File A. The two ATAC-seq datasets have rather low numbers of uniquely mapped reads. Especially the ATAC1 dataset has only ∼ 5% uniquely mapped reads. Hence, we decided to use only DNase and ATAC2 for bench-marking.

For both, the DNase and the ATAC2 dataset we used JAMM (34) for peak calling with parameters “-f 1 -d y”, which results in JAMM only considering 5’ ends of reads and retaining duplicate reads.

We used the open-source library Jstacs (35, 36) (class `projects.encodedream.Pileup`) to calculate the 5’ coverage with ATAC-seq or DNase-seq reads at each position and normalized coverage relative to the mean of a 10000 bp sliding window.

### Model

The statistical model behind EpiTALE is based on modelling the total binding score of a putative target box ***x*** = *x*_0_*x*_1_ … *x*_*L*_ to the RVD sequence ***r*** = *r*_1_*r*_2_ … *r*_*L*_ of length *L*. Each RVD *r*_*l*_ ∈ {*AA*,…, *Y Y, A*\*,…,*Y*\*} is composed of its two amino acids. Each putative target box can be a sequence of *x*_ℓ_ ∈ {*A, C, G, T*}and *x*_0_ denotes the nucleotide at position 0 of the target box.

In addition to the original definition of the PrediTALE model, we introduce *q*_ℓ_ ∈ [0, 1] as the methylation level at position ℓ and ***q*** ∈ *q*_1_ … *q*_*L*_ as the sequence of methylation levels for each nucleotide of the target box ***x*** bound by an RVD.

The total binding score of a putative target box ***x*** given the RVD sequence ***r*** of a TALE and the methylation probabilities ***q*** is the sum of the following terms: i) The dependency between the zero-th nucleotide and the first RVD, ii) binding between first RVD and first nucleotide, and iii) binding of the remaining RVDs to the remaining nucleotides, where ii) and iii) may be weighted by a position-dependent but sequence-independent term.

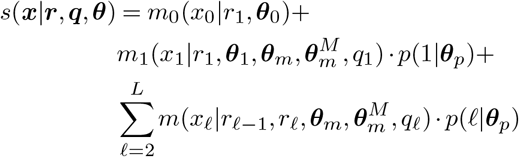

The set of real-valued parameters 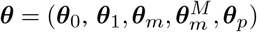 includes the terms for binding to the zero-th, first and remaining nucleotide, the binding specifities for 5mC as well as the position-dependent term.

As in the original PrediTALE model, the term for binding to the zero-th nucleotide *m*_0_(*x*_0_ | *r*_1_, ***θ***_0_) is independent of methylation levels, since there are no appropriate activity or binding studies regarding methylation sensitivity available, yet. As in PrediTALE, this term corresponds to the sum of the following parameter values: i) the *a-priori* parameter of nucleotide zero 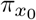, ii) the parameter 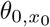 for the zero-th nucleotide and in case that *r*_1_ is in set ℛ_0_ iii) the parameter 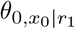 for the zero-th nucleotide depending on *r*_1_, where *ℛ*_0_ = {*HD, NN, NG, NI, NS*} and *π*_*T*_ = log(0.6), *π*_*C*_ = log(0.3), *π*_*A*_ = *π*_*G*_ = log(0.05).

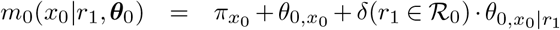

The binding of the first RVD to the first nucleotide of the target box is modelled by the term 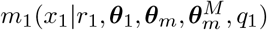 and consists of the sum of the following two main terms: i) The probability (1 − *q*_1_) that the first position is unmethylated is multiplied by the term adopted from the original PrediTALE model. ii) Given a methylation level *q*_1_ *>* 0 at position 1, the methylation level is multiplied by the preference of the 13th AA of the first RVD to bind to a methylated cytosine. If the first RVD *r*_1_ belongs to the set ℛ_1_, the general preference of the complete first RVD to a methylated cytosine is added.

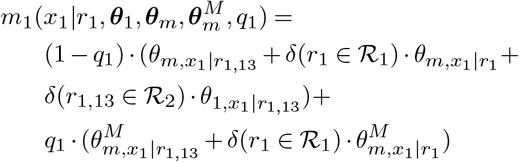

The sets are set to ℛ_1_ = {*HD, NN, NG, HG, NI, NK*} and ℛ_2_ = {*D, N, G, I*} as originally proposed for PrediTALE (13).

The binding to the remaining positions is modelled by terms 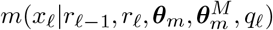, which are identical to the previous PrediTALE variant in the non-methylated case. In case of a methylation level greater than 0 at position ℓ, the preference of the 13th amino acid to bind a methylated cytosine and, if applicable, the preference of the entire RVD for a methylated cytosine is included.

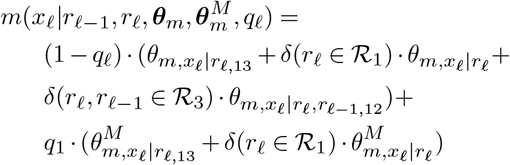

In analogy to the original PrediTALE publication, we set ℛ_3_ = {*HD, NN, NG, NI*}

We set *q*_ℓ_ := 0, if the nucleotide at position *l* of the target sequence is not a cytosine.

### Scale parameters to model specificities for 5mC

As described in the previous section, we extended the previously trained PrediTALE model by adding parameters for the specificity to bind to ‘5mC’ to incorporate methylation information into the TALE target prediction of EpiTALE. The former training (13) included pairs of TALEs and their putative target boxes from different experiments (37–41).

A thorough study by Zhang *et al*. tested all theoretically possible combinations of RVDs to bind to 5methylcytosine (5mC), 5-hydroxymethylcytosine (5hmC), cytosine and thymine (16). For this purpose, the activation of a GFP expression reporter was measured in the screening. To obtain fitted values for the parameters 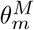 (see above) representing binding preferences to methylated cytosines, we considered specificities for methylated cytosines determined by Zhang *et al*. (16). To this end, we scaled the measured values from Zhang *et al*. to match the range of parameter values of the original PrediTALE model. Specifically, we used the two reference points for cytosine and thymine also present in the data of Zhang *et al*. to scale the raw measured values 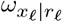 to fit to our trained parameter space.

Let 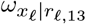 be the arithmetic mean of all 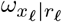 with the same 13th AA.

The specificity of the 13th AA of the RVD to bind 5*mC* is determined by the following scaling:

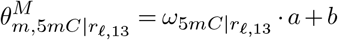

with

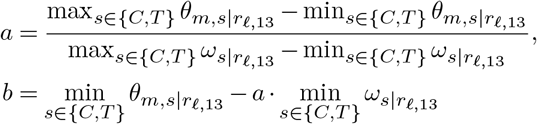

The parameters for the specificity of RVDs from ℛ_1_ to bind 5*mC* result from the following scaling:

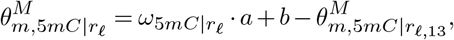

with

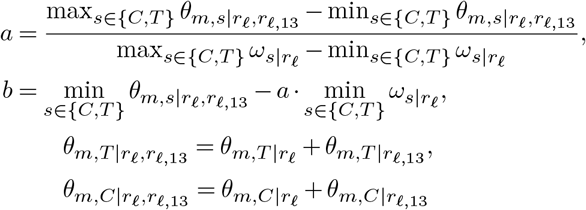

Here, we consider only values measured for 5*mC* but not those measured for 5*hmC* for two reasons. First, the plant genome contains much less 5*hmC* than 5*mC*. Second, it is not possible to distinguish between 5mC and 5hmC in bisulfite sequencing. We also decided not to calculate the average over both measurements, since the specificity of both differs substantially for some RVDs.

A visualization of the model parameters of EpiTALE including the parameters for 5mc is shown in Figure 1. Several of the thirteenth amino acids and several of the common RVDs show large differences in specificity between an unmethylated and a methylated cytosine.

**Fig. 1.**
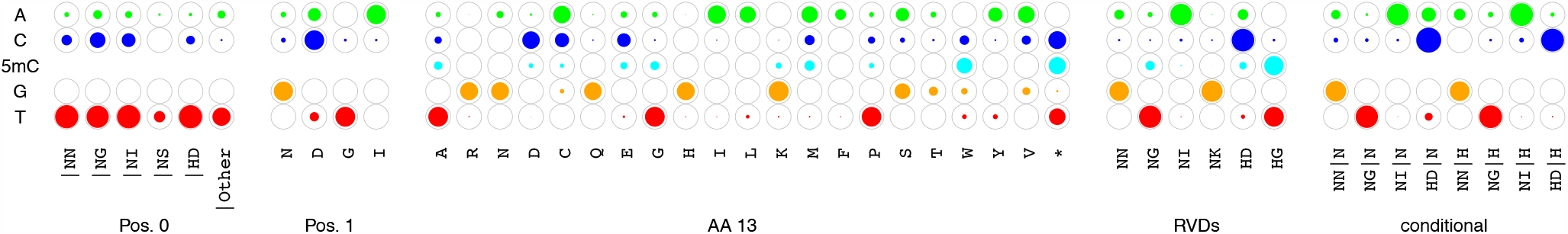
Parameters of the EpiTALE model represented by circles filled to a degree proportional to specificity parameters. There are no separate parameters for methylated cytosines for the sub-model at position 0, position 1 and the “conditional” sub-model, as there is no sufficient data available yet.

### Accessibility filter

Target boxes predicted by EpiTALE may further be filtered for chromatin accessibility. For each predicted target sequence, a window from 300 bp upstream to 50 bp down-stream of the target box is checked for an overlapping peak within the peaks determined by JAMM (34) from chromatin accessibility data.

As an additional filter criterion, the number of positions that correspond to at least one 5’-end of a read within a defined region around the predicted target box is considered. For protomerome-wide predictions, this region corresponds to the complete promoter sequence (−300 bp to +200 bp relative to TSS (12)). For genome-wide prediction, a window around the predicted target box (−300 bp to +200 bp) is considered. If there is an overlapping peak *or* the coverage filter criterion is fulfilled, we consider the target box as accessible.

### Prediction of TALE target boxes

The basic procedure of score calculation in a sliding window along the input sequences remains as described previously (13). Additionally, we use the scaled parameters for methylation specificity and the methylation levels from WGBS-seq data for promoterome-wide TALE target prediction. Here, we compare promoterome-wide predictions with and without methylation information and we perform these studies for each TALE of 3 *Xoo* strains and 10 *Xoc* strains (Supplementary File F).

In addition, DNase-seq and ATAC-seq data were used to check the predicted targets for accessibility and to derive a filter criterion based on the predictions for TALEs from the 3 *Xoo* strains. We then applied this fixed filter criterion to the predictions for TALEs from the 10 *Xoc* strains.

### Genome-wide predictions & filtering

With EpiTALE, we perform genome-wide predictions in the *Oryza sativa* Nipponbare genome (MSU7) including methylation information and the filter based on chromatin accessibility. For the resulting top 100 predictions of each TALE, we use above mentioned RNA-seq data to search for a differentially expressed region near the putative target box using DerTALE, as described previously (13). We visualize the resulting profiles with an auxiliary R script, which plots the RNA-seq profile surrounding the putative target box and uses gff3 files to display known genes overlapping with the profile. Here, we use the MSU7 annotation (http://rice.plantbiology.msu.edu/pub/data/Eukaryotic_Projects/o_sativa/annotation_dbs/pseudomolecules/version_7.0/all.dir/all.gff3). For differentially expressed regions with no overlapping annotated gene, we use blastx and blastn to search for similar sequences in the non-redundant protein sequence (nr) database and the reference RNA sequences (refseq_rna) database using NCBI BLAST+ version 2.7.1 (42) (ftp://ftp.ncbi.nlm.nih.gov/blast/executables/blast+/LATEST/).

### Evaluation of prediction results

To compare the impact of the two epigenetic features, we evaluate the following prediction variants: original PrediTALE model without epigenetic features (P), EpiTALE with consideration of methylation (P + Methyl), EpiTALE with filtering based on the accessibility filter criterion (P + Access) and EpiTALE with methylation and accessibility filtering (P + Methyl + Access).

We compare the performance of these 4 variants for the above mentioned *Xoo* and *Xoc* strains based on the corresponding RNA-seq infection studies. For benchmarking based on differentially expressed genes, we consider a promoter region 300 bp upstream of the transcription start site to 200 bp down-stream of the transcription start site or until the start codon as described previously (12, 13).

The use of RNA-Seq data from inoculation studies to evaluate the predictions entails two problems: First, when plant tissue is inoculated with a *Xanthomonas* strain, multiple TALEs lead to differential gene expression. Hence, it is not possible to clearly assign differentially expressed genes to a particular TALE. The RVD sequences of the TALEs of the Xanthomonas strains studied are given in F. Secondly, it is not clear whether a gene was up-regulated by the binding of a TALE to its promoter or by secondary effects triggered through inoculation with the *Xanthomonas* strain. So, we define all genes as true positive (TP) target genes that are upregulated in the RNA-seq data relative to control and have a predicted target box within the promoter. We define those genes as false positives (FP) that are not up-regulated after inoculation, but have a predicted target box in their promoter. The definition of false negatives is not clearly possible, since up-regulated genes without a predicted target box could be indirect target genes.

In order to compare the 4 EpiTALE prediction variants mentioned above, we proceed in analogy to the previous comparison of PrediTALE with alternative approaches (13): We vary the number *t* of predictions per TALE considered between 1 and 50, i.e., we only consider the *t* predictions with the largest prediction scores for each TALE. Within these top lists, we determine the number of TPs for each cutoff.

### Availability

The EpiTALE suite is available as a JavaFX-based standalone application with graphical user interface and as command line application under http://jstacs.de/index.php/EpiTALE. A minimal example for testing is available from zenodo at https://www.doi.org/10.5281/zenodo.4749294. Source code is available from https://github.com/Jstacs/Jstacs in package `projects.tals.epigenetic`.

The EpiTALE suite contains tools, that (i) convert BedMethyl files to Bismark format, (ii) merge two Bismark files, (iii) compute a coverage pileup of 5’ ends of mapped reads from an DNase-seq or ATAC-seq experiment, (iv) normalize the coverage pileup relative to the mean of a 10000 bp sliding window, (v) convert methylation data (Bismark files) and chromatin accessibility data (coverage pileup and/or narrow-Peak file) to promoter coordinates and (vi) predict TALE target boxes with optional epigenetic input within genomic or promoter sequences.

## RESULTS/DISCUSSION

### Impact of epigenetic features on individual target boxes

In this section, we illustrate the potential impact of considering epigenetic features on TALE target prediction before turning to a more systematic evaluation in the subsequent sections. Specifically, we consider individual examples of TALE target boxes that were shown to be affected either by DNA methylation or by chromatin accessibility and demonstrate that, for these examples, TALE target prediction would improve when considering epigenetic features.

#### Accessibility of TALE target boxes

As an additional way of improving the selection of target boxes predicted by EpiTALE, we implemented the option to provide chromatin accessibility data for filtering the predicted target boxes. Basically, this corresponds to using chromatin accessibility data to remove inaccessible target boxes from the list of predictions.

In Figure 2, we compare the accessibility of two almost identical predicted targets boxes, which only differ in one nucleotide. These are two predicted target boxes of TalAE15 from *Xoo* ICMP3125^*T*^. In panel (A), one predicted target box is located in the promoter of gene Os04g39400. Among the predictions for TalAE15, this target box appears on rank 17 using PrediTALE or EpiTALE without epigenetic features. However, in all three accessibility datasets this putative target box appears to be inaccessible and the downstream gene it is not up-regulated after infection with *Xoo* ICMP3125^*T*^ in the RNA-seq experiment. Hence, this target box appears to be a false positive prediction, which could be removed from the list of predictions when filtering for accessibility as implemented in EpiTALE.

**Fig. 2.**
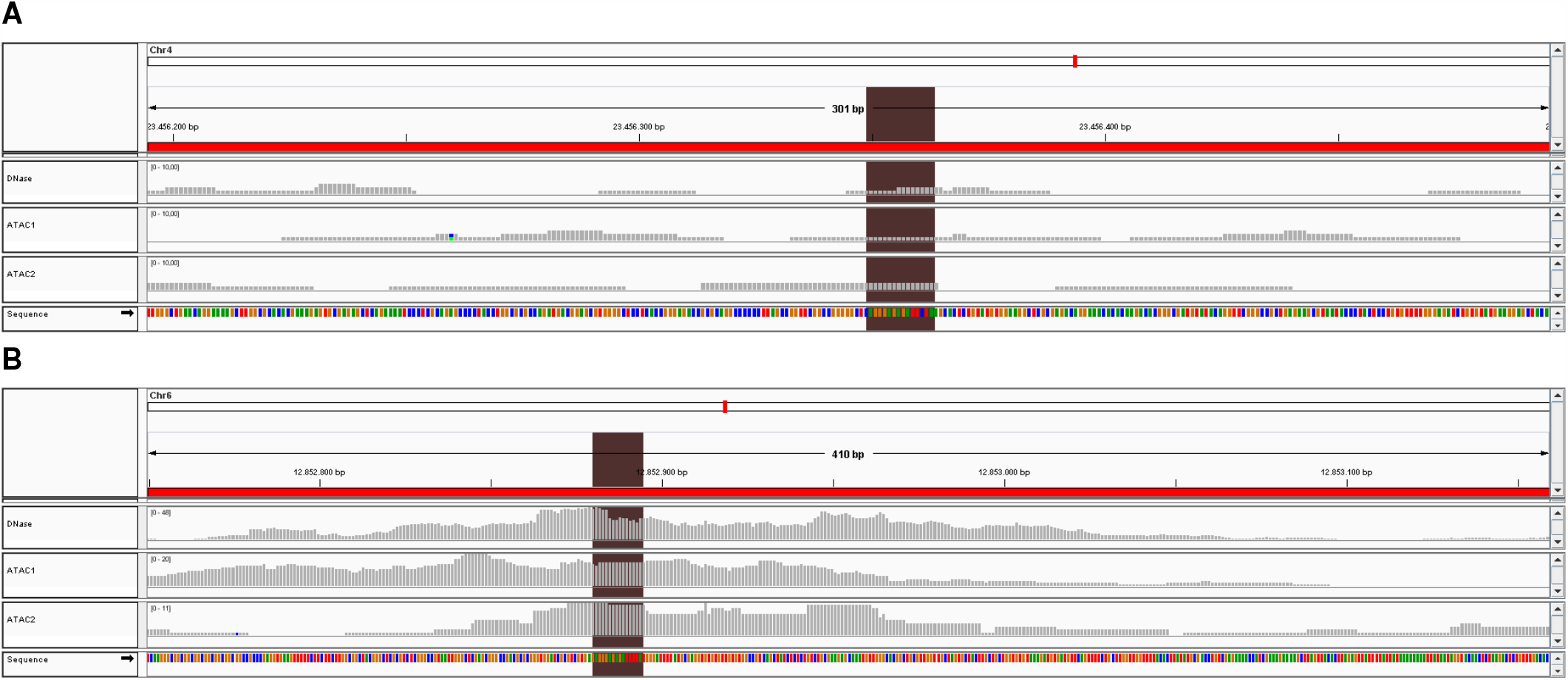
IGV (43) snapshot comparing promotor accessibility based on DNAse-seq and ATAC-seq data of two almost identical target boxes predicted by PrediTALE. (A) Coverage profile of the Os04g39400 promoter region. (B) Coverage profile of the Os06g22140 promoter region. The brown box marks the predicted target box.

The almost identical putative target box from Figure 2 (B) receives rank 54 among the TalAE15 predictions with standard PrediTALE and is located in the promoter of gene Os06g22140. The IGV snapshot shows high accessibility in all 3 accessibility datasets within and around the box. The coverage in the DNase dataset is approximately 2.5 and 5 times as high as in the ATAC1 and ATAC2 datasets, respectively. Furthermore, the downstream gene is up-regulated according to RNA-seq experiments, and we would consider it a true positive prediction. Given that putative false positive predictions as shown in Figure 2A would be filtered for being inaccessible, well-accessible target boxes like that in the promoter of Os06g22140 would appear on better ranks in the list of EpiTALE predictions.

#### Methylation of TALE target boxes

It has been shown previously (15, 16) that the specificity of RVDs for methylated cytosines differs from unmethylated cytosines. With the extension of PrediTALE to include separate specificities for methylated cytosines (cf. Methods), we allow for considering methylation levels along the input sequences in TALE target prediction.

An example of a methylated putative target site is shown in Figure 3. According to the WGBS-seq data described in the Methods section, the promoter of Os01g57140 shows methylation at several positions. Here, methylation is also present in the predicted target box of TalAB16 from *Xoo* ICMP3125^*T*^. Without paying attention to methylation, this target box appears on rank 19 within the PrediTALE predictions for TalAB16, whereas we know from RNA-seq data after infection that this is not a true positive target. If we provide methylation data to EpiTALE, this target box receives a substantially lower score and ends up on rank 208 of the predictions. Hence, including methylation information resulted in removing a putative false positive prediction from the topranking predictions of EpiTALE.

**Fig. 3.**
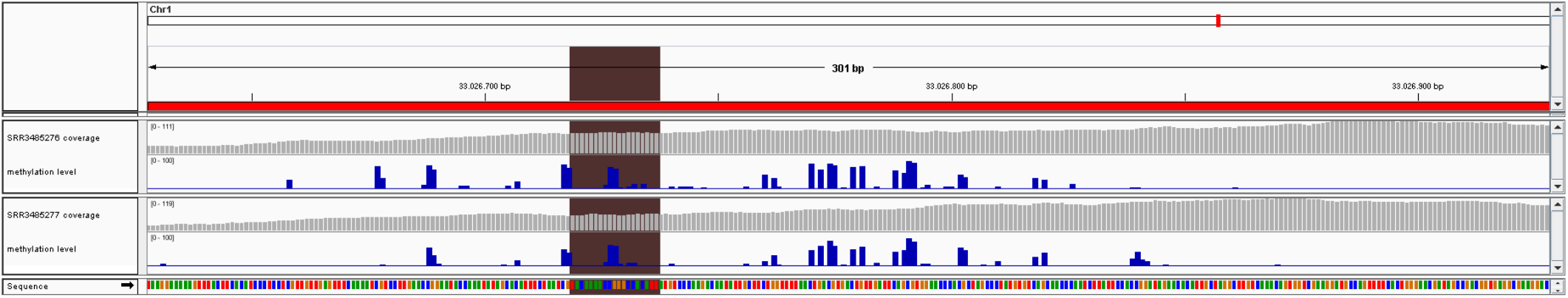
IGV (43) snapshot with BS-seq coverage track and methylation level track in the vicinity of a predicted target boxes in rice. The brown box marks the predicted target box of TalAB16 on the promoter of gene Os01g57140.

### Performance evaluation of different accessibility filter parameters on DNase dataset

In this section, we benchmark the effect of different filters based on chromatin accessibility applied to EpiTALE predictions. To this end, we test different filtering thresholds for predicting target boxes of the TALEs present in 3 *Xoo* strains, and we evaluate the chosen filter criteria on independent data for 10 *Xoc* strains.

Supplementary Figure S2 shows violin plots of the accessibility values for the three accessibility datasets considered. Here, we compare the accessibility of true positive (TP) EpiTALE predictions compared with false positive (FP) EpiTALE predictions according to RNA-seq data. EpiTALE predictions are generated for TALEs present in *Xoo* and the *Xoc* strains and chromatin accessibility is summarized per predicted target box as the average normalized coverage around the target box. When averaging, we always consider the window starting 300 bp upstream and ending 50 bp downstream the target box in the strand orientation of the downstream gene.

The violin plot of the DNase dataset shows a visible although small difference in accessibility between TP and FP targets for both *Xoo* and *Xoc*. The two ATAC-seq datasets, however, show substantially smaller differences with almost identical median values and generally low coverage. These ATAC-seq datasets are likely of limited use for filtering TALE target predictions. Hence, we focus on the DNase-seq data in the following analyses, and provide results using the ATAC2 dataset as supplementary figures. The reason that the two ATAC-seq datasets are less suited for filtering TALE target predictions may be the relatively low genomic coverage with ATAC-seq reads but also different experimental conditions when collecting these publicly available ATAC-seq data. Plants at different life stages and grown under different greenhouse conditions may have different accessibility profiles. For the DNase-seq data, this issue seems to be less severe. We speculate that ATAC-seq data collected under the same conditions as for the infection experiments might still be informative for TALE target prediction.

The accessibility filter criterion consists of two parts: A putative target box from the initial predictions survives the filter if it has an overlapping peak of chromatin accessibility within the window from 300 bp upstream to 50 bp downstream of the target box. A putative target box also survives the filter if at least *t* positions within the complete promoter show coverage greater than zero.

The performance of EpiTALE using different thresholds *t* for filtering based on the DNase-seq dataset compared with the original PrediTALE neglecting chromatin accessibility is shown in Supplementary Figure S3 for three *Xoo* strains. Here, the number of true positive (TP) target boxes is plotted against the number of predictions allowed per TALE, with a rank cutoff from 1 to 50. To ensure comparability to the original PrediTALE publication (13), we use the same type of performance plots with the same rank cutoffs and the same definition for differentially up-regulated genes caused by the respective strains in the RNA-seq infection studies. Briefly, we consider those genes as putatively up-regulated by TALEs that have an uncorrected p-value below 0.05 in the RNA-seq infection studies of the 3 Xoo strains and are at least 2-fold up-regulated.

Using the accessibility filter, a threshold of 30 yields the largest area under the curve (AUC) of TP predictions for the strains ICMP 3125^T^ and PXO83. In case of ICMP 3125^T^, filtering with this threshold for any rank cutoff shows improved or at least identical performance as PrediTALE without filtering, and a larger improvement than any other filter threshold tested. For PX083, the same effect can be observed, where only when considering the top 3 predictions per TALE, a threshold of 35 results in one additional TP. For PXO142, accessibility filtering with a threshold of 30 within a range of 1 to 30 predictions per TALE increases or at least retains the number of TP predictions. For higher rank cutoffs, the filtering results in one TP less than for the unfiltered version. Thus, a threshold of 30 leads to a reduction of TPs only in rare cases and mostly leads to an increase in the number of TPs within the top predictions. In complete analogy, supplementary Figure S7 shows performance plots for the 10 *Xoc* strains. For 8 of the 10 strains, the accessibility filtering based on the DNase dataset results in a higher AUC for each of the 6 thresholds. However, for 7 of these 8 strains a threshold of 25 results in the highest AUC. The threshold of 30 chosen from the *Xoo* datasets does not result in the optimal result for *Xoc*, but performs substantially better than the original PrediTALE version without filtering. Filtering based on the DNase-seq dataset works slightly worse only for strains CFBP7331 and CFBP7341.

For the predictions of the top 50 target boxes of each TALE of the *Xoo* and *Xoc* strains considered, the proportion of TP and FP target boxes that pass the accessibility filter is shown in supplementary Figure S9. The TP target boxes are usually accessible according to the accessibility filter criterion with a threshold of 30. FP target boxes in turn are rather filtered out as they are occasionally inaccessible.

### Performance of EpiTALE model considering epigenetic DNA modifications

In this section, we further investigate the effect of including methylation-specific parameters into the EpiTALE model, and its combination with the accessibility filter studied in the previous section. Specifically, we consider four modelling alternatives: i) the original PrediTALE model (P), ii) the EpiTALE model including specificities for methylated cytosines (P + Methyl), iii) the PrediTALE model combined with the accessibility filter (P + Access), and iv) the EpiTALE model combined with the accessibility filter (P + Methyl + Access).

The results of the performance evaluation of these four alternatives for TALE target prediction of *Xoo* TALEs are shown in Figure 4. Here, accessibility is determined based on the DNase dataset. The number of true positive predictions is improved by either of the epigenetic features for the strains ICMP 3125^T^ and PXO83, where the improvement due to the accessibility filter is more pronounced than the improvement due to including methylation levels into the EpiTALE model. For both strains, performance is further increased by combining both epigenetic features. For PXO142, the accessibility filter alone leads to slightly decreased prediction performance, whereas methylation information alone as well as the combination of both epigenetic features leads to a slight improvement compared with the original PrediTALE variant.

**Fig. 4.**
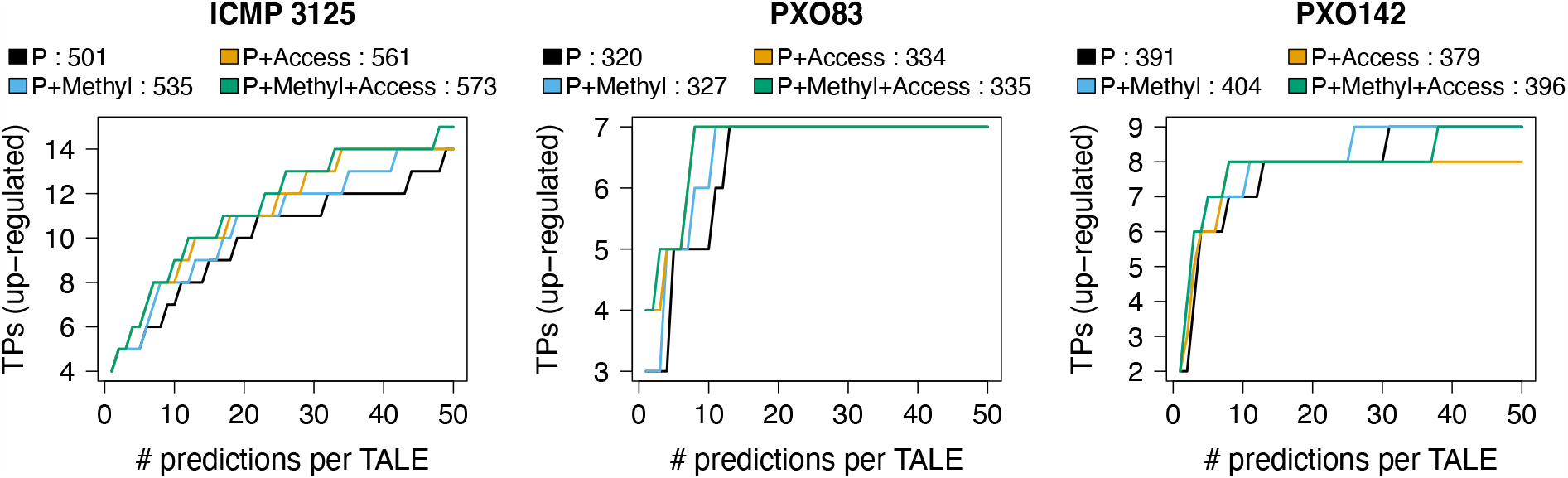
EpiTALE performance evaluation for three *Xoo* strains considering epigenetic features. We plot the number of predicted target genes that are also up-regulated in the infection (true positives, TPs) against the number of predicted target sites per TALE for PrediTALE (P) and three EpiTALE variants including only methylation information (P+Methyl), only filtering based on chromatin accessibility (P+Access), or a combination of both (P+Methyl+Access). In the legends, we further report the area under the curve for PrediTALE and the individual EpiTALE variants.

The results for the *Xoc* strains are shown in Figure 5. For 8 of the 10 strains, methylation information, filtering according to target box accessibility, and the combination of both epigenetic features lead to a clear increase of AUC compared to PrediTALE without these features. For CFBP7331 and CFBP7341, only considering the methylation information leads to an improvement, because the accessibility criterion for these two strains is too strict in some cases and a few TP target boxes are determined to be inaccessible.

**Fig. 5.**
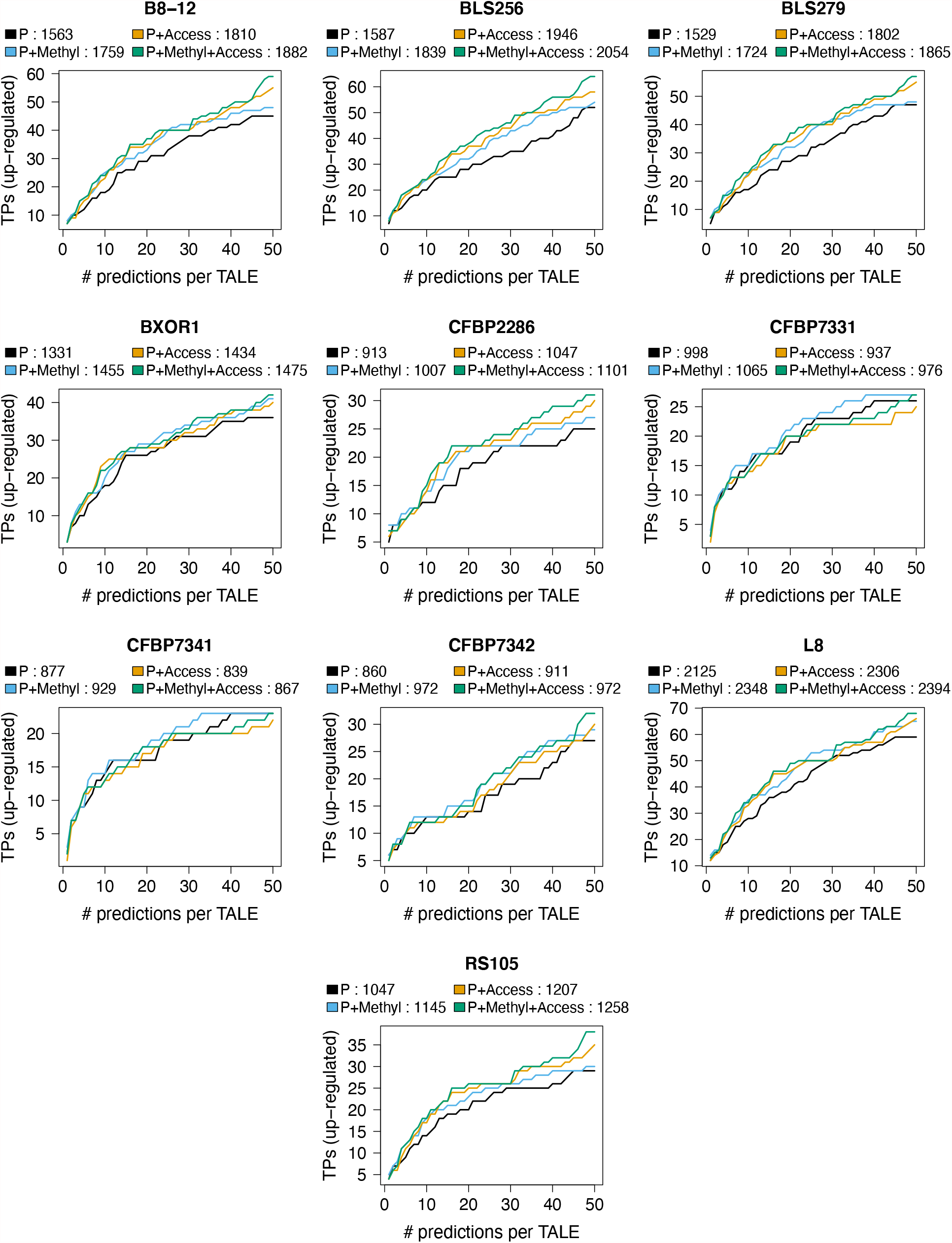
EpiTALE performance evaluation for ten *Xoc* strains considering epigenetic features. We plot the number of predicted target genes that are also up-regulated in the infection (true positives, TPs) against the number of predicted target sites per TALE for PrediTALE (P) and three EpiTALE variants including only methylation information (P+Methyl), only filtering based on chromatin accessibility (P+Access), or a combination of both (P+Methyl+Access). In the legends, we further report the area under the curve for PrediTALE and the individual EpiTALE variants.

The performance based on the accessibility dataset ATAC2 for 3 *Xoo* strains is presented in Supplementary Figure S5. In this case, the performance of the EpiTALE variants for which accessibility is used for filtering is substantially lower. In order to rule out the possibility that the decrease in performance is simply due to the chosen filtering criteria, we tested different thresholds for this dataset as presented in Figure S6. However, none of the thresholds considered leads to restoring the performance of the original PrediTALE variant.

### Considering epigenetic features improves ranks of true positive targets

In this section, we focus on the top 20 predictions of the four EpiTALE variants for three *Xoo* and ten *Xoc* strains that also show upregulation after infection with the respective strains. The complete list of true positive predictions is given in Supplementary Table C, and the subset of predictions for *Xoo* strains is provided in Table 1. For each of the 3 *Xoo* strains, all three EpiTALE variants including epigenetic features mostly yield an improvement of the rank of the true positive target gene compared with the original PrediTALE variant without epigenetic features. The strongest rank improvement is almost always achieved by the EpiTALE variant that considers methylation of the target box as well as its accessibility. However, an improvement can often be observed already when considering only one of the epigenetic features. The gene Os09g07460, coding for a kelch repeat protein, is among the top 20 predictions for TalBA8 for all three EpiTALE variants considering epigenetic features. This gene has not been among the top 20 predictions of the original Predi-TALE variant, but has been reported by Talvez (11, 13). Regarding target boxes predicted for TALEs from ten *Xoc* strains (cf. Supplementary Table C), all three EpiTALE variants including epigenetic features in the majority of cases either yield an improved or an unchanged prediction rank for true positive genes.

**Table 1.** Putative TALE target genes that are among the top 20 predictions per TALE for any of the four approaches. For each *Xoo* strain, we list the gene ID (MSU7) and the log fold change (lfc) in the corresponding RNA-seq experiment. For each of the four EpiTALE variants, we further list the TALE(s), for which a gene has been predicted as a target and in parentheses the corresponding prediction rank.

| Gene | lfc | P | P+Methyl | P+Access | P+Methyl+Access | annotation |
| --- | --- | --- | --- | --- | --- | --- |
| **ICMP 3125<sup>T</sup>** |  |  |  |  |  |  |
| Os02g06670 | 3.815 | TalBA8 (1) | TalBA8 (1) | TalBA8 (1) | TalBA8 (1) | retrotransposon protein |
| Os09g29820 | 2.819 | TalAR13 (2) | TalAR13 (2) | TalAR13 (2) | TalAR13 (2) | OsTFX1 - bZIP transcription factor |
| Os03g51760 | 2.734 | TalAD22 (9) | TalAD22 (7) | TalAD22 (6) | TalAD22 (6) | OsFBX109 - F-box protein |
| Os04g05050 | 2.221 | TalAB16 (11) | TalAB16 (8) | TalAB16 (7); | TalAB16 (7) | pectate lyase |
| Os01g40290 | 1.894 | TalAA15 (1) | TalAA15 (1) | TalAA15 (1) | TalAA15 (1) | expressed protein |
| Os05g45070 | 1.704 | TalAO15 (15) | TalAO15 (13) | TalAO15 (11) | TalAO15 (10) | harpin-induced protein 1 |
| Os11g26790 | 1.695 | TalAH11 (1) | TalAH11 (1) | TalAH11 (1) | TalAH11 (1) | dehydrin |
| Os06g03710 | 1.591 | TalES1 (19) | TalES1 (17) | TalES1 (13) | TalES1 (12) | DELLA protein SLR1 |
| Os01g73890 | 1.079 | TalBM2 (1) | TalBM2 (1) | TalBM2 (1) | TalBM2 (1) | transcription initiation factor IIA gamma |
| Os10g28240 | 0.918 | TalAR13 (6) | TalAR13 (6) | TalAR13 (4) | TalAR13 (4) | calcium-transporting ATPase |
| Os09g07460 | 0.746 | TalBA8 (22) | TalBA8 (19) | TalBA8 (18) | TalBA8 (17) | kelch repeat protein |
| **PXO142** |  |  |  |  |  |  |
| Os02g49350 | 5.163 | TalBH2 (8) | TalBH2 (5) | TalBH2 (7) | TalBH2 (5) | plastocyanin-like |
| Os03g09150 | 2.530 | TalBH2 (4) | TalBH2 (3) | TalBH2 (4) | TalBH2 (3) | pumilio-family RNA binding |
| Os11g31190 | 2.514 | TalBH2 (3) | TalBH2 (2) | TalBH2 (3) | TalBH2 (2) | SWEET14 (nodulin MtN3) |
| Os09g29820 | 2.272 | TalAR14 (3) | TalAR14 (2) | TalAR14 (2) | TalAR14 (2) | OsTFX1 - bZIP transcription factor |
| Os03g51760 | 1.368 | TalAD23 (13) | TalAD23 (11) | TalAD23 (8) | TalAD23 (8) | OsFBX109 - F-box protein |
| Os01g40290 | 0.887 | TalAA16 (1) | TalAA16 (1) | TalAA16 (1) | TalAA16 (1) | expressed protein |
| Os06g29790 | 0.833 | TalAO16 (4) | TalAO16 (4) | TalAO16 (3) | TalAO16 (3) | phosphate transporter 1 |
| Os07g06970 | 0.824 | TalAP15 (1) | TalAP15 (1) | TalAP15 (1) | TalAP15 (1) | HEN1 |
| **PXO83** |  |  |  |  |  |  |
| Os09g29820 | 2.82 | TalAR3 (5) | TalAR3 (4) | TalAR3 (1) | TalAR3 (1) | OsTFX1 - bZIP transcription factor |
| Os02g06670 | 2.74 | TalAR3 (83);<br>TalBA2 (1) | TalAR3 (74);<br>TalBA2 (1) | TalAR3 (52);<br>TalBA2 (1) | TalAR3 (48);<br>TalBA2 (1) | retrotransposon protein |
| Os03g51760 | 1.91 | TalAD5 (13) | TalAD5 (11) | TalAD5 (8) | TalAD5 (8) | OsFBX109 - F-box protein |
| Os04g19960 | 1.70 | TalAC5 (1) | TalAC5 (1) | TalAC5 (1) | TalAC5 (1) | retrotransposon protein |
| Os04g05050 | 1.62 | TalAB5 (11) | TalAB5 (8) | TalAB5 (7) | TalAB5 (7) | pectate lyase |
| Os07g06970 | 1.40 | TalAP3 (1) | TalAP3 (1) | TalAP3 (1) | TalAP3 (1) | HEN1 |
| Os03g03034 | 1.18 | TalAQ3 (5) | TalAQ3 (4) | TalAQ3 (4) | TalAB5 (97);<br>TalAQ3 (3) | flavonol synthase |

The accessibility filter criterion appears to be inappropriate for some putative target boxes upstream of true positive target genes, which are determined to be inaccessible, although they are upregulated in the RNA-seq experiments. This applies to the putative target box in the promoter of Os01g52130 for TalBF members from *Xoc* strains B8-12, BLS256, BLS279, CFBP2286, BXOR1, CFBP7331, CFPB7341, CFPB7342, L8, RS105; the putative target box upstream of Os02g06130 for TalAF from B8-12 and L8; the putative target box upstream of Os07g01490 for TalBD from B8-12, BLS256, BLS279, BXOR1, L8; the putative target box upstream of Os07g03279 for TalBE from BXOR1, CFBP7331, CFPB7341; the putative target box upstream of Os03g22020 for TalBU from CFBP7331 and the putative target box upstream of Os12g06930 for TalBI from CFPB7342. Out of 323 true positive target boxes, the majority of 212 target boxes, however, obtains an improved rank when considering both epigenetic features, while the rank of 87 target boxes remains unchanges compared with the original Predi-TALE variant. Among the target genes with an improved prediction rank are well known TALE targets like Os07g06970 coding for HEN1, but also promising novel candidates like Os03g53800 a beta-glucosidase precursor.

Both the methylation information and chromatin accessibility considered in this study have been derived from publicly available datasets that have been collected for different purposes and scientific questions. Hence, these have been determined under different conditions, e.g., from different plants at a different life stage than for the infection studies that are represented by the RNA-seq data. On the one hand, this may explain both the lowered ranks of the above-mentioned true positive target genes when considering methylation information, but also target genes that are up-regulated in the infection studies not passing the accessibility filter. On the other hand, the widely improved prediction ranks for many of the remaining true positive target genes provide a strong indication that both types of data provide valuable information for TALE target prediction. Our results suggest that with matched WGBS-seq and DNase-seq/ATAC-seq data of sufficient quality, the quality of computational TALE target predictions could be boosted even further.

### Genome-wide TALE target prediction considering DNA methylation and chromatin accessibility

Independently of existing gene annotations, we performed genome-wide predictions of TALE target boxes in *Oryza sativa* Nipponbare (MSU7) for 3 *Xoo* and 10 *Xoc* strains using the EpiTALE version with methylation and DNase accessibility data. As accessibility filter criterion, we select regions around the binding site that are similar to the promoter setting. Specifically, a peak within chromatin accessibility data must be present in a region from -300 bp to +50 bp relative to the target box or at least 30 positions within a window of -300 bp to +200 bp relative to the target box must correspond to the 5’-end of a DNase-seq read.

To determine differentially expressed regions near predicted target boxes, we use our tool DerTALE (13) and the mapped RNA-Seq from above-mentioned infection studies. For DerTALE we use the same settings as in the original PrediTALE publication (13). Briefly, we search for differentially expressed regions of at least 300 bp within a region of *±*3000 bp around the top 100 predicted target boxes of each TALE. Genome-wide prediction shows that for 16 *Xoo* TALEs, differential expressed regions are close to at least one predicted target box. In total, we obtain 20 of such target boxes (complete list in Supplementary Table D), of which 13 have also been observed in the previous prediction limited to promoters. Among these 20 target boxes, 18 have already been reported in the original PrediTALE publication (13). By using the two epigenetic features in the EpiTALE variant, we obtain 2 novel target boxes near differentially expressed regions. Figure 6 presents the RNA-seq profile in the region of a target box predicted for members of family TalAB on chromosome 2. The putative target boxes of TalAB5 (PXO83) and TalAB16 (ICMP 3125^T^) are identical and do not overlap with a gene annotation known from MSU7. We extracted the sequences under the differentially expressed regions, and first compared them against the NCBI protein database ‘nr’ using blastx but received no matching result. We additionally compared these sequences against the NCBI reference RNA sequences (‘refseq_rna’) using blastn, which resulted in a predicted mRNA, coding for a calcium-transporting ATPase (XM_015770644.2).

**Fig. 6.**
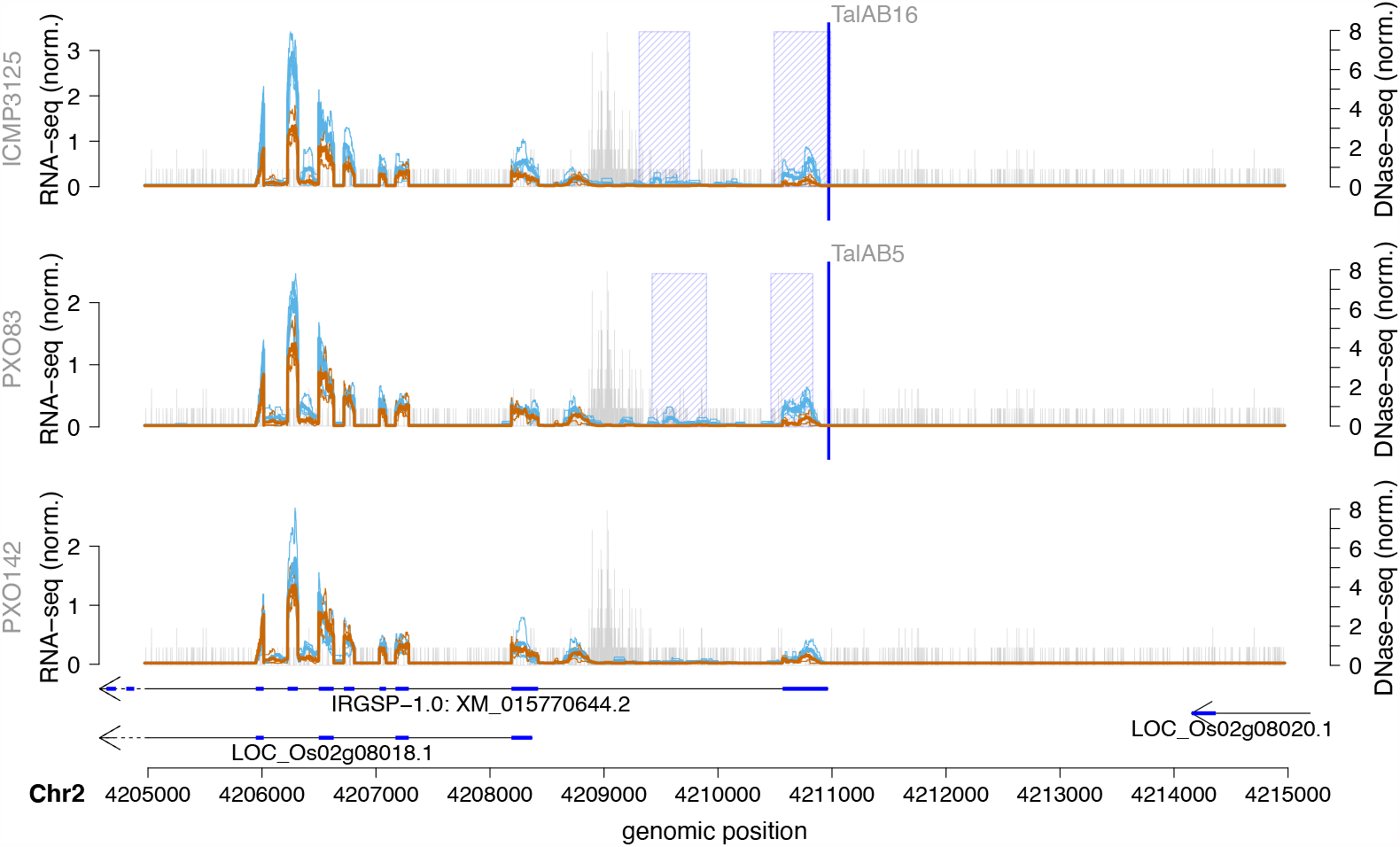
Genome-wide predictions of TalAB in *Oryza sativa* Nipponbare profile for 3 *Xoo* strains in the area of the TalAB target box. RNA-seq coverage after inoculation (blue line) is compared with mock control (brown line). In addition, we show the average of individual replicates of control and treatment are summarized as thick lines. The blue shaded boxes mark the differentially expressed regions. The arrows under the profiles reflect the MSU7 annotation within the genomic region. The genomic position of the TALE target box is marked by a vertical blue line. Vertical grey bars indicate the number and 5’-position of reads in the DNase data.

However, one putative target box reported from genome-wide predictions in the original PrediTALE publication (Os04g05050) appears on a lower rank due to methylation of the target box, which might be caused by non-matching experimental conditions as discussed previously.

A complete list of the genome-wide predictions of TALEs from the ten *Xoc* strains is given in Supplementary Table E, of which we select two examples for a detailed discussion. The first of these is a predicted target box of members of the TalAX family located on chromosome 1. Members of TalAX are present in all ten *Xoc* strains. The corresponding class tree is shown in Supplementary Figure S11 and the RNA-seq profile around the putative target box is provided in Figure 7. Close to the putative target box is a differentially expressed region that has no overlapping MSU7 gene annotation. For 7 of the 10 strains, the predicted target box is among the top 100 predictions. For strains CFBP7331 and CFPB7341, this target box appears only in the top 200 prediction due to differences in the RVD sequence of the TalAX members present in these strains. However, the RNA-seq data suggest that these TALEs are still capable of activating downstream expression since a differential region is detected for these strains as well. TalAX2 from CFBP7342 deviates even further from the RVD composition of the remaining strains, and no target box in this region was predicted for TalAX2. In agreement with this prediction, we do not observe a differentially expressed region after *Xoc* CFBP7342 infection. For the sequences under this differentially expressed region, database search using blastx and blastn against ‘nr’ and ‘refseq_rna’, respectively, did not result in a match.

**Fig. 7.**
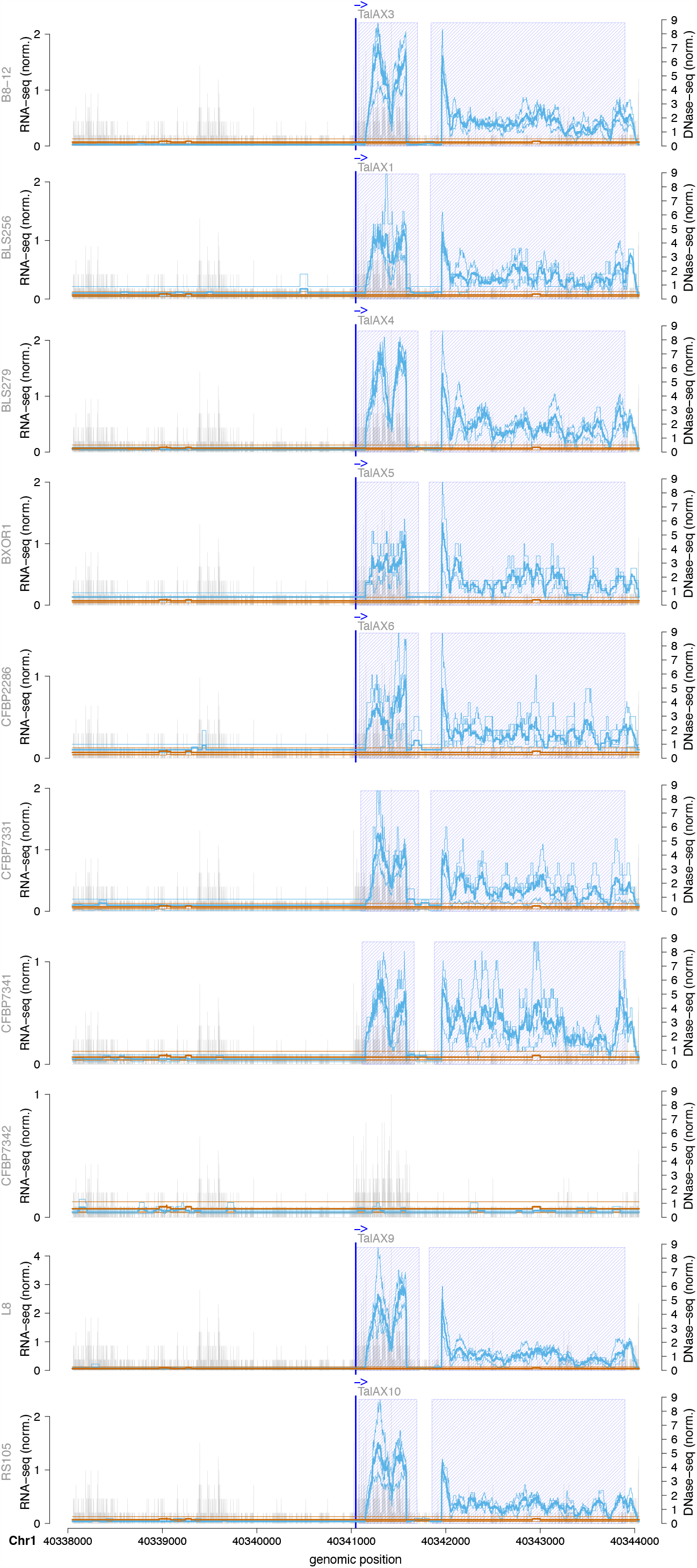
Genome-wide predictions of TalAX in *Oryza sativa* Nipponbare profile for 10 *Xoc* strains in the area of the TalAX target box. RNA-seq coverage after inoculation (blue line) is compared with mock control (brown line). In addition, we show the average of individual replicates of control and treatment are summarized as thick lines. The blue shaded boxes mark the differentially expressed regions. The genomic position of the TALE target box is marked by a vertical blue line. Vertical grey bars indicate the number and 5’-position of reads in the DNase data.

As a second example, we discuss a putative target box on chromosome 6 for members of the TalBN class present in 8 of 10 *Xoc* strains. The corresponding class tree is shown in Supplementary Figure S12 and the RNA-seq profile around the binding site is presented in Figure 8. The TalBN members from 7 of the 8 strains have identical RVD sequences, whereas TalBN2 of CFBP7342 show one difference in RVD sequence. This target box on chromosome 6 is among the top 100 predictions only for these 7 TalBN members and DerTALE report a differentially expressed region after infection with these strains. The remaining TalBN members (TalBN2 of CFBP7342) has no putative target box among the top 100 predictions at this position, and the region shows no differential expression as well as for the 2 strains with no TalBN member (CFBP7341, CFBP7331). This indicates that this differentially expressed region may be caused by TalBN members of the strains with the putative target site. The differentially expressed region does not overlap with an annotated MSU7 gene and the corresponding sequence had no matches in BLAST searches.

**Fig. 8.**
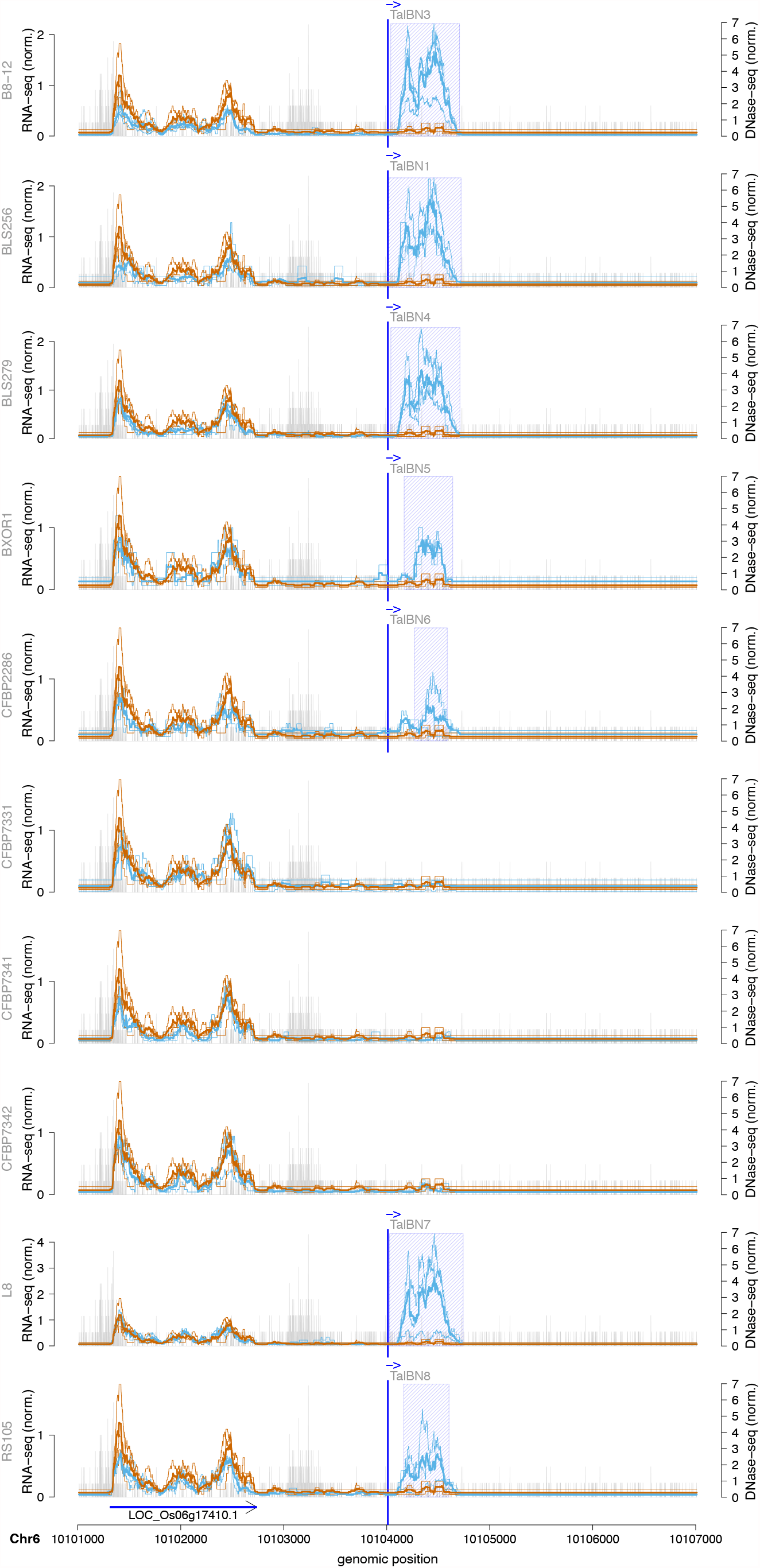
Genome-wide predictions of TalBN in *Oryza sativa* Nipponbare profile for 10 *Xoc* strains in the area of the TalBN target box. RNA-seq coverage after inoculation (blue line) is compared with mock control (brown line). In addition, we show the average of individual replicates of control and treatment are summarized as thick lines. The blue shaded boxes mark the differentially expressed regions. The arrows under the profiles reflect the MSU7 annotation within the genomic region. The genomic position of the TALE target box is marked by a vertical blue line. Vertical grey bars indicate the number and 5’-position of reads in the DNase data.

## Conclusion

With the goal of improving the prediction of TALE targets, we present EpiTALE, an extended version of PrediTALE including epigenetic features. Both, methylation levels and the chromatin accessibility around putative target sites have a decisive impact on the likelihood of being bound by a TALE. Even if a putative target box matches the specificity of the RVDs of a TALE, inaccessibility of the respective chromatin may inhibit binding and thus inhibit activation of the transcription of the downstream gene (44, 45). We demonstrate that for the prediction of TALE target boxes, the consideration of the epigenetic state of rice plants leads to an improved quality of TALE target predictions by EpiTALE. For many true positive *Xoo* and *Xoc* target boxes, EpiTALE yielded improved prediction ranks of true positive targets compared with the original PrediTALE variant. Nevertheless, there are still false positive predictions and we suggest an experimental verification of novel targets.

We perform promoter and genome-wide predictions and find several predictions common to both approaches, but we also find target boxes upstream of differentially expressed regions in RNA-seq infection studies that do not overlap with a currently annotated gene.

The use of the epigenetic features is optional for the user. Depending on the availability of data, only methylation and/or chromatin accessibility data can be provided to EpiTALE to improve target prediction. In our study, the strongest improvement in accuracy was achieved by considering both epi-genetic features in EpiTALE. Our results suggest that collecting condition-matched WGBS-seq and DNase-seq/ATAC-seq data may further improve the quality of computational TALE target predictions. The EpiTALE suite presented here provides the means necessary to integrate such data into TALE target prediction and is available from http://jstacs.de/index.php/EpiTALE.

## Supporting information

Supplementary Table A. Mapping statistics of ATAC and DNase-seq data.

Supplementary Table B. List of positive and negative targets for Xoo and Xoc.

Supplementary Table C. Complete list of top 20 predictions for all four varaiants and Xoo and Xoc strains.

Supplementary Table D. Genome-wide prediction for three Xoo strains.

Supplementary Table E. Genome-wide prediction for ten Xoc strains.

Supplementary File F. RVD sequences of all Xoo and Xoc TALEs considered in this manuscript.

## FUNDING

This work was supported by grants from the Deutsche Forschungsgemeinschaft (http://www.dfg.de) (BO 768 1496/8-1 to JB and GR 4587/1-1 to JG) and CA16107 “EuroXanth” (https://euroxanth.eu) to JB. The funders had no role in study design, data collection and analysis, decision to publish, or preparation of the manuscript.

This preprint is formatted using a L^A^T_E_X class by Ricardo Henriques

