## Supplementary Figures S1 - S12, Captions of Supplementary Tables A - E and Supplementary File F for "Epigenetic features improve TALE target prediction"

### SUPPLEMENTARY MATERIAL

#### Supplementary Figures

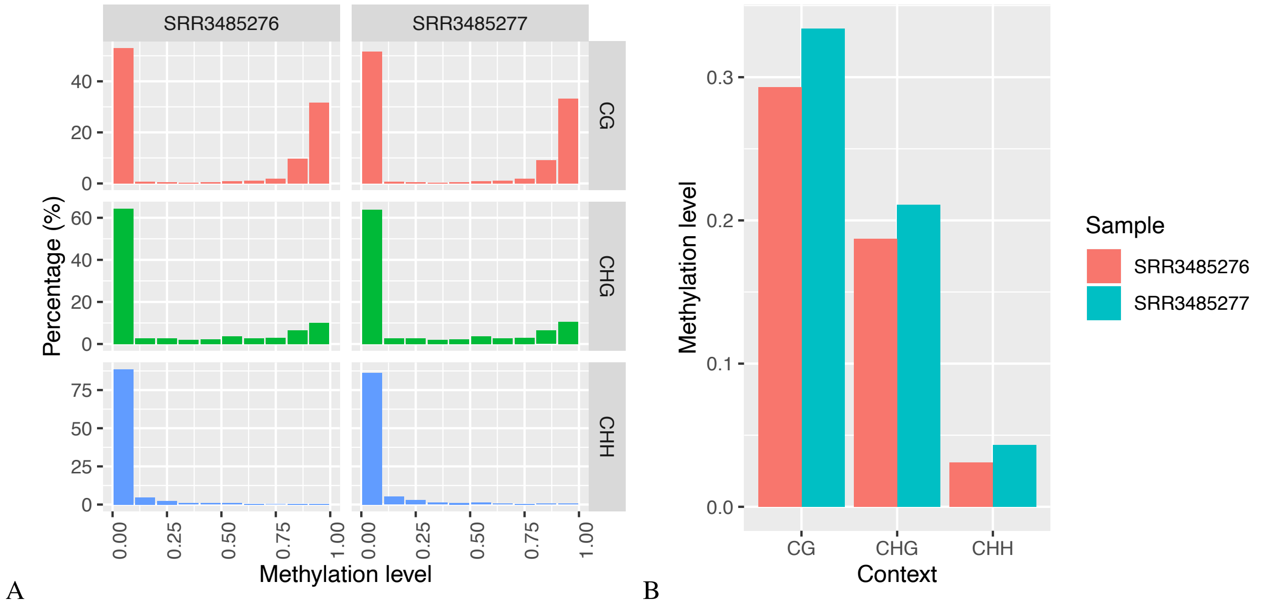


**Supplementary Figure S1.** ViewBS (28) visualization of BS-seq data of rice from European Nucleotide Archive run accession SRR3485276 and SRR3485277 with (A) distribution of methylation level and (B) global methylation level for three contexts.

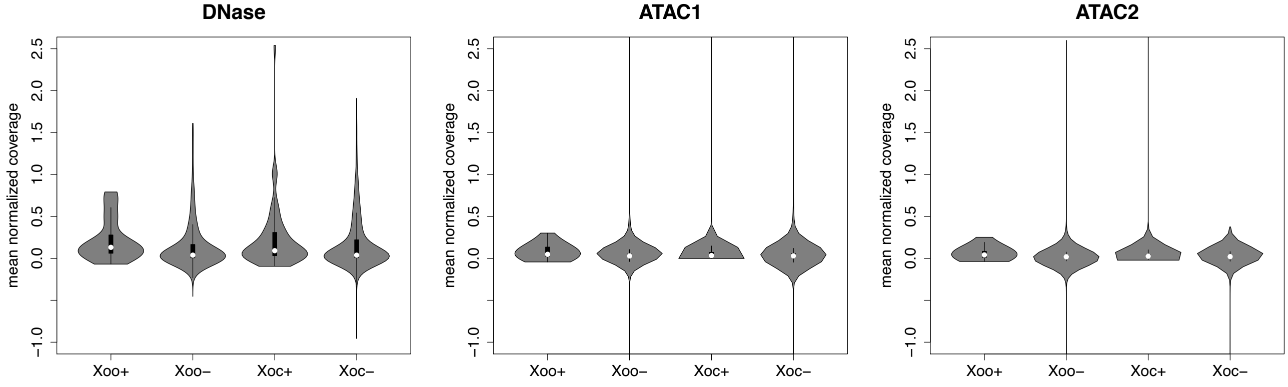


**Supplementary Figure S2.** The chromatin accessibility in the region of positive (+) and negative (-) target sites as violin plots of mean normalized coverage for three chromatin accessibility datasets separately for *Xoo* and *Xoc* strains.

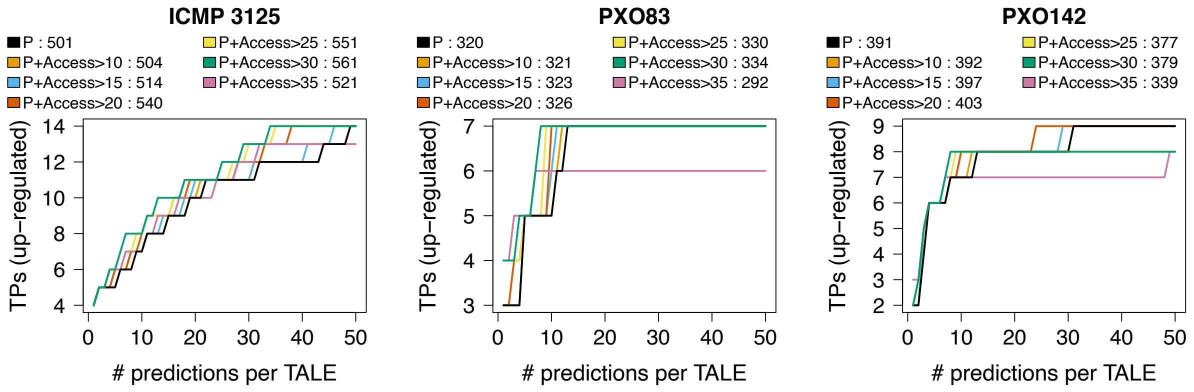


**Supplementary Figure S3.** EpiTALE performance evaluation for three *Xoo* strains considering the accessibility of the promoter of the target gene using DNase-seq data based on the number of predicted target genes that are also up-regulated in the infection (true positives, TPs) against the number of predicted target sites per TALE. EpiTALE without filtering is compared with different threshold for the filter criterion. The area under the curve for different criteria is shown above the diagram.

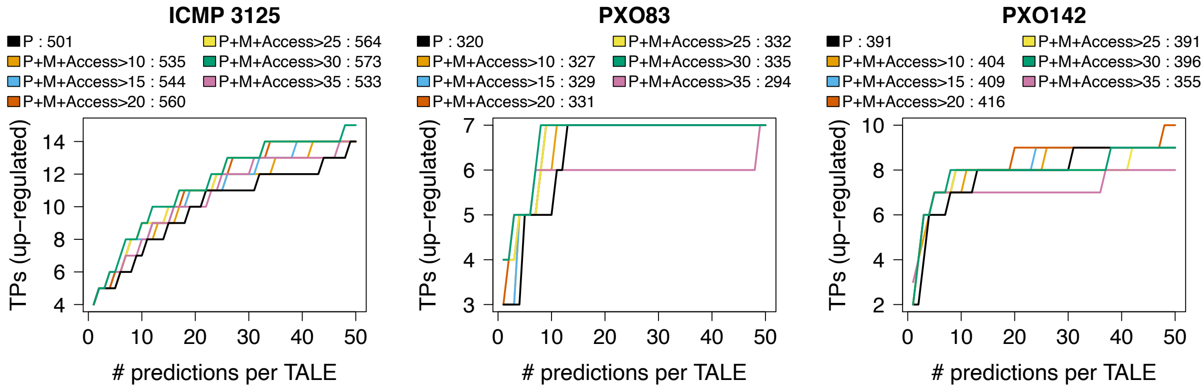


**Supplementary Figure S4.** EpiTALE performance evaluation for three *Xoo* strains considering the accessibility of the promoter of the target gene using DNase-seq data based on the number of predicted target genes that are also up-regulated in the infection (true positives, TPs) against the number of predicted target sites per TALE. EpiTALE without filtering and without attention on methylation is compared with different threshold for the filter criterion on predictions, where methylation is considered. The area under the curve for different criteria is shown above the diagram.

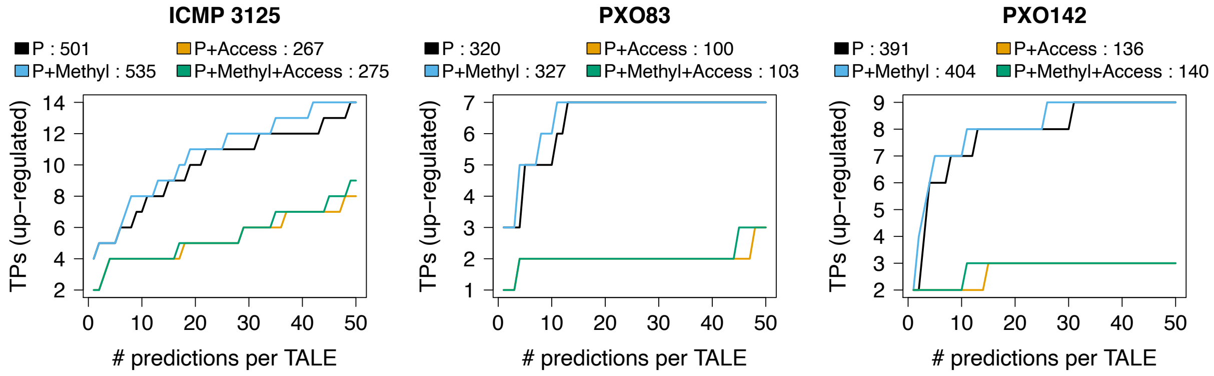


**Supplementary Figure S5.** EpiTALE performance evaluation for three *Xoo* strains considering the accessibility of the promoter of the target gene using ATAC2 dataset based on the number of predicted target genes that are also up-regulated in the infection (true positives, TPs) against the number of predicted target sites per TALE. The 4 mentioned EpiTALE variants are compared. The area under the curve for different criteria is shown above the diagram.

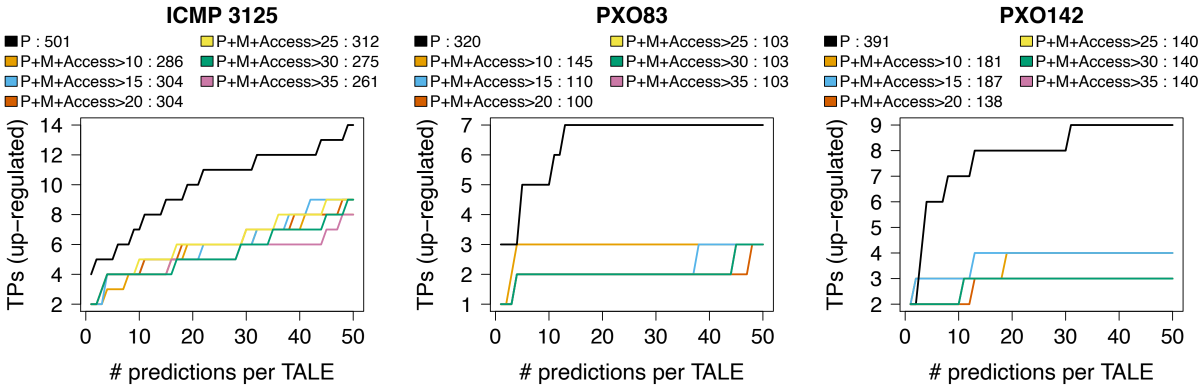


**Supplementary Figure S6.** EpiTALE performance evaluation for three *Xoo* strains considering the accessibility of the promoter of the target gene using ATAC-seq data based on the number of predicted target genes that are also up-regulated in the infection (true positives, TPs) against the number of predicted target sites per TALE. EpiTALE without filtering is compared with different threshold for the filter criterion and considering the methylation. The area under the curve for different criteria is shown above the diagram.

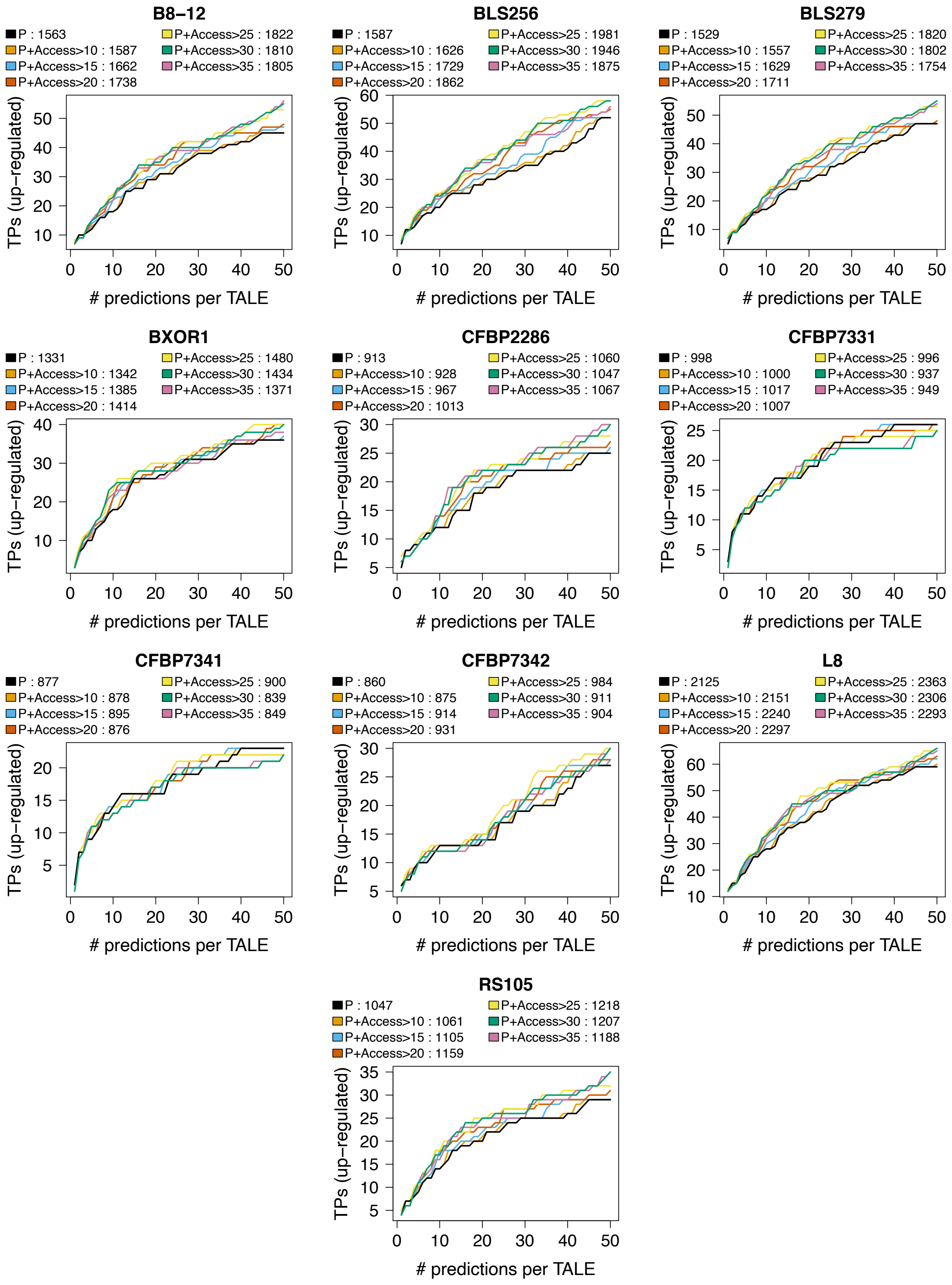


**Supplementary Figure S7.** EpiTALE performance evaluation for ten *Xoc* strains considering the accessibility of the promoter of the target gene using DNase-seq data based on the number of predicted target genes that are also up-regulated in the infection (true positives, TPs) against the number of predicted target sites per TALE. EpiTALE without filtering is compared with different threshold for the filter criterion. The area under the curve for different criteria is shown above the diagram.

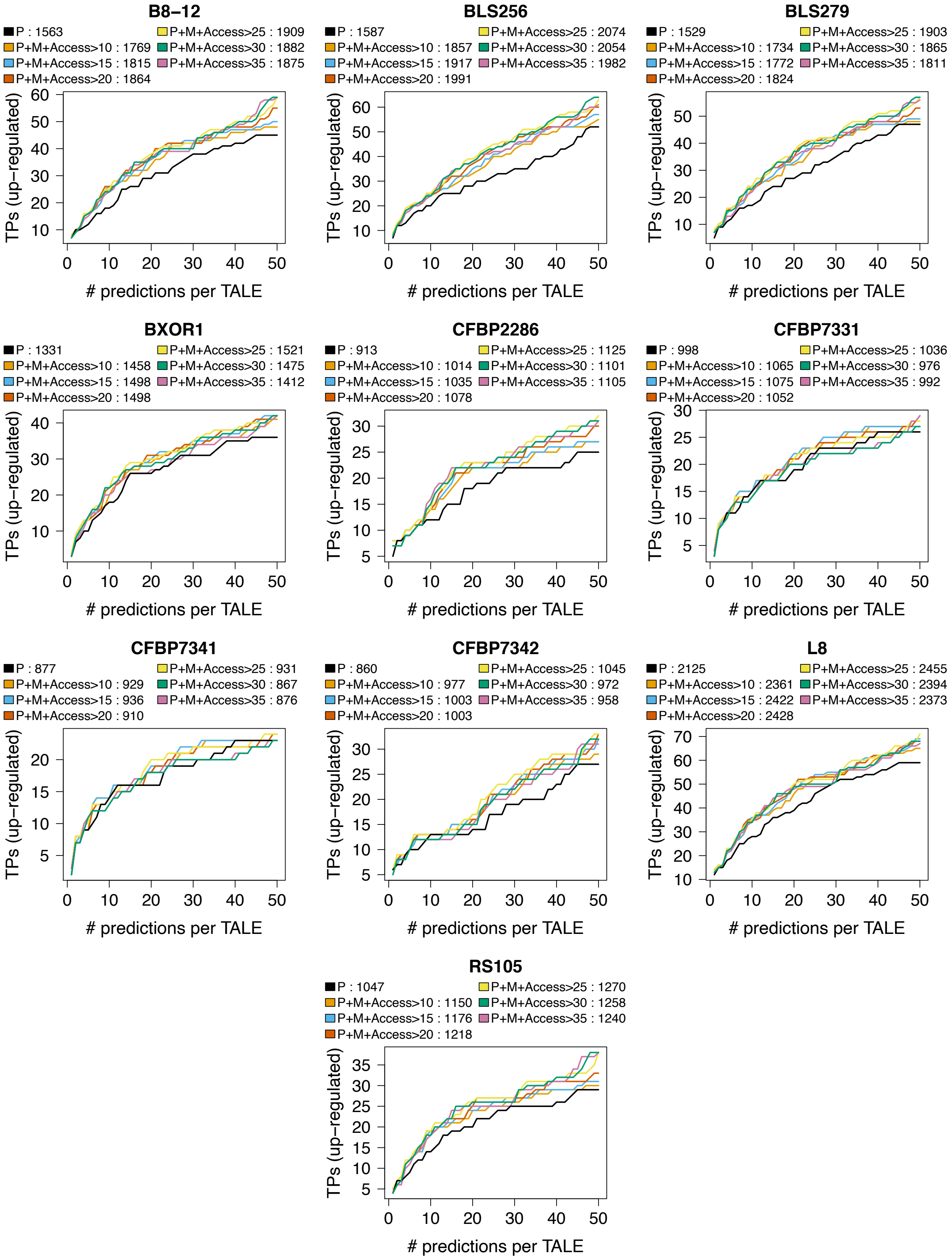


**Supplementary Figure S8.** EpiTALE performance evaluation for ten *Xoc* strains considering the accessibility of the promoter of the target gene using DNase-seq data based on the number of predicted target genes that are also up-regulated in the infection (true positives, TPs) against the number of predicted target sites per TALE. EpiTALE without filtering and without attention on methylation is compared with different threshold for the filter criterion on predictions, where methylation is considered. The area under the curve for different criteria is shown above the diagram.

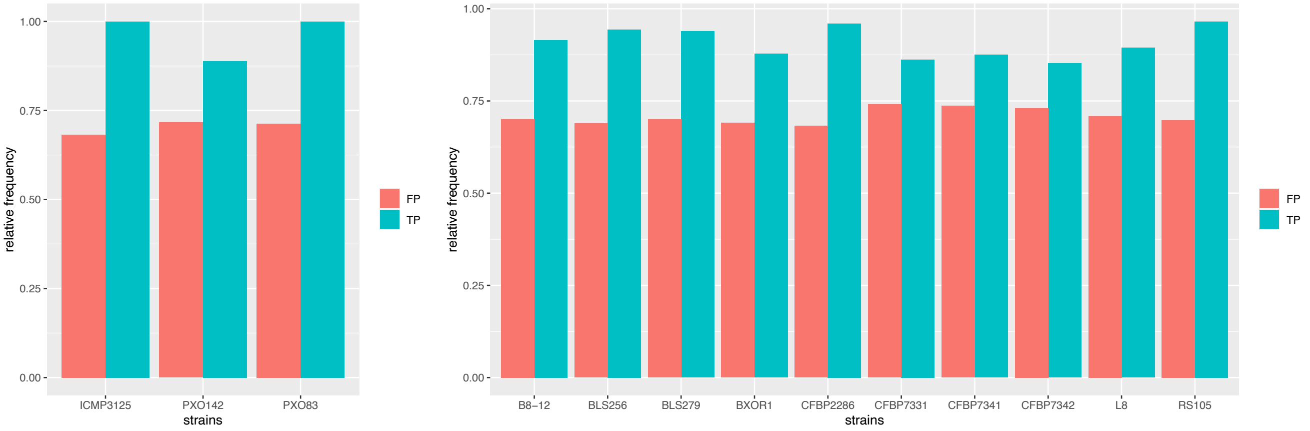


**Supplementary Figure S9.** Proportion of true positive (TP) and false positive (FP) prediction, that survive filter criterion, within top 50 predictions of each TALE.

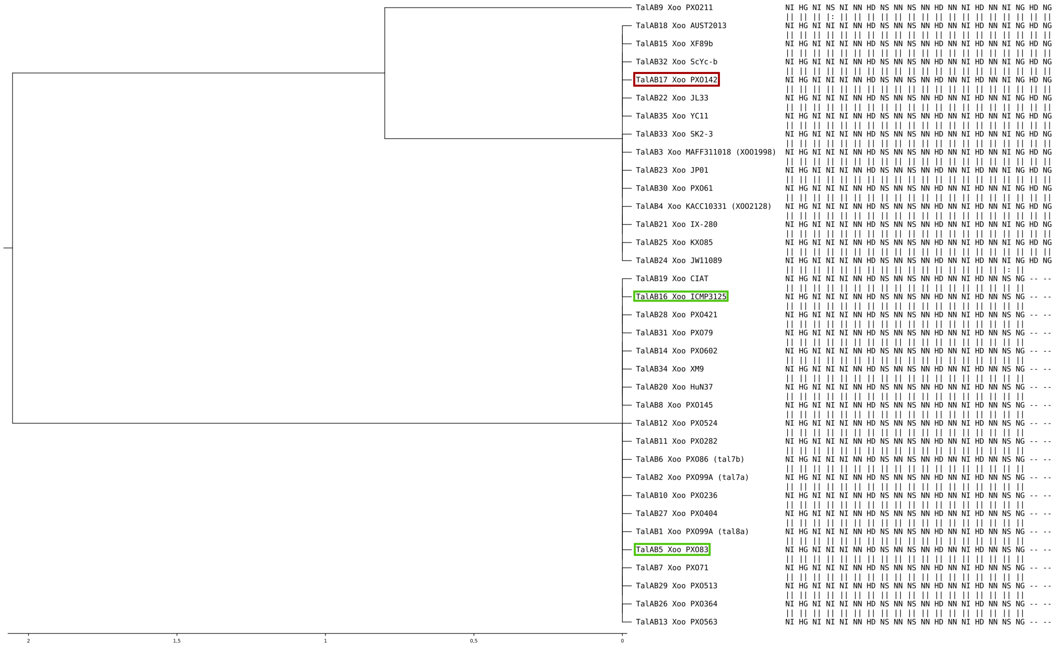

**Supplementary Figure S10.** AnnoTALE (46) class tree for TalAB. Class members are marked in green if the corresponding target box of Figure 6 is within the top 100 predictions of EpiTALE with both epigenetic features and a differentially expressed region according to RNA-seq infection studies is in the surrounding area. If this target box does not occur within the top 500 predictions, the member is marked in red.

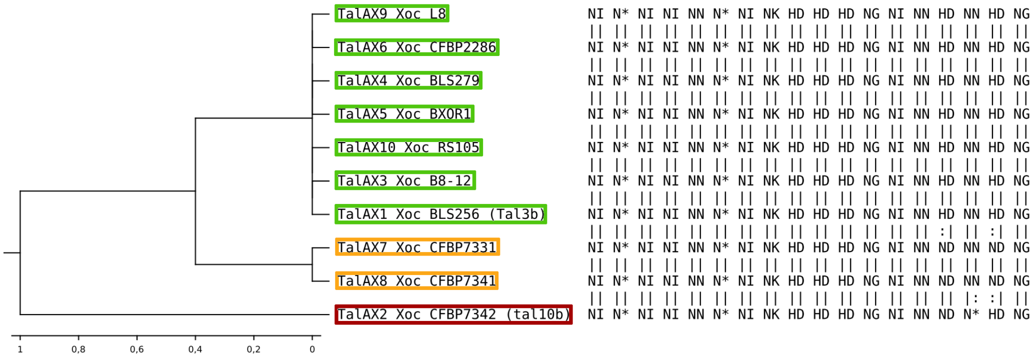


**Supplementary Figure S11.** AnnoTALE (46) class tree for TalAX. Class members are marked in green if the corresponding target box of figure 7 is within the top 100 predictions of EpiTALE with both epigenetic features and a differentially expressed region according to RNA-seq infection studies is in the surrounding area. They are marked in orange if the target box is within the top 200 predictions and has a respective differentially expressed region. If this target box does not occur within the top 500 predictions, the member is marked in red.

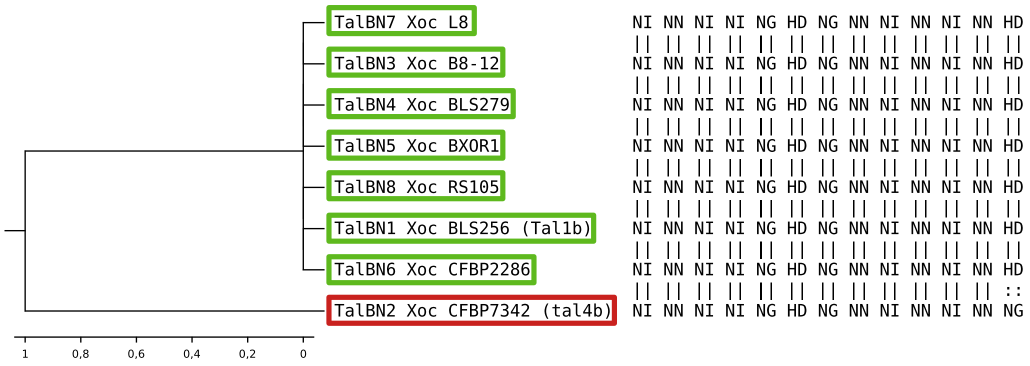


**Supplementary Figure S12.** AnnoTALE (46) class tree for TalBN. Class members are marked in green if the corresponding target box of figure 8 is within the top 100 predictions of EpiTALE with both epigenetic features and a differentially expressed region according to RNA-seq infection studies is in the surrounding area. If this target box does not occur within the top 500 predictions, the member is marked in red.

#### Supplementary Tables

**Supplementary Table A.** Mapping statistics of ATAC and DNase-seq data.

[Available as separate XLS file `dnase_atac_seq_statistics_mapped.xls`]

**Supplementary Table B.** List of positive and negative targets for *Xoo* and *Xoc*.

[Available as separate XLS file `positives_negatives_Xoo_Xoc.xls`]

**Supplementary Table C.** Complete list of top 20 predictions for all four varaiants and *Xoo* and *Xoc* strains.

[Available as separate XLS file `predictions4variants.xls`]

**Supplementary Table D.** Genome-wide prediction for three *Xoo* strains.

[Available as separate XLS file `summaryGenomewideXoo.xls`]

**Supplementary Table E.** Genome-wide prediction for ten *Xoc* strains.

[Available as separate XLS file `summaryGenomewideXoc.xls`]

**Supplementary File F.** RVD sequences of all *Xoo* and *Xoc* TALEs considered in this manuscript.

[Available as separate FastA file `TALE_RVDs.fasta`]
